# DeepSV: Accurate calling of genomic deletions from high-throughput sequencing data using deep convolutional neural network

**DOI:** 10.1101/561357

**Authors:** Lei Cai, Yufeng Wu, Jingyang Gao

## Abstract

**Background:** Calling genetic variations from sequence reads is an important problem in genomics. There are many existing methods for calling various types of variations. Recently, Google developed a method for calling single nucleotide polymorphisms (SNPs) based on deep learning. Their method visualizes sequence reads in the forms of images. These images are then used to train a deep neural network model, which is used to call SNPs. This raises a research question: can deep learning be used to call more complex genetic variations such as structural variations (SVs) from sequence data?

**Results:** In this paper, we extend this high-level approach to the problem of calling structural variations. We present DeepSV, an approach based on deep learning for calling long deletions from sequence reads. DeepSV is based on a novel method of visualizing sequence reads. The visualization is designed to capture multiple sources of information in the sequence data that are relevant to long deletions. DeepSV also implements techniques for working with noisy training data. DeepSV trains a model from the visualized sequence reads and calls deletions based on this model. We demonstrate that DeepSV outperforms existing methods in terms of accuracy and efficiency of deletion calling on the data from the 1000 Genomes Project.

**Conclutions:** Our work shows that deep learning can potentially lead to effective calling of different types of genetic variations that are complex than SNPs.

**Availability and implementation:** DeepSV’s source code and sample result as part of this project are readily available from GitHub at https://github.com/CSuperlei/DeepSV/.

## 1 Introduction

High-throughput DNA sequencing technologies have generated vast amount of sequence data. These data enable novel approaches for studying many important biological questions. One example is calling genetic variations such as single nucleotide polymorphisms (SNPs) or structural variations (SVs) from sequence data. There have been many existing computational methods for sequence-based calling of SNPs or SVs. For example, for SNP calling, one popular caller is GATK [1]. On the high level, calling genetic variations from sequence data can be viewed as a classification problem in machine learning. That is, given the sequence data at a candidate variant site, we are to classify the site into one of the two categories: variant or wild-type. Among many existing classification approaches, deep learning based on e.g. convolutional neural network (CNN) is becoming increasingly popular. CNN has outperformed existing approaches in a number of important applications. Among these, the most noticeable application of CNN is image processing, where deep learning has significantly improved the state of the art [2]. A natural research direction is using CNN for genetic variant calling with sequence data. Recently, Google’s DeepVariant [3] was developed to call SNPs and short insertion/deletions (indels) from sequence data. The key idea of DeepVariant is viewing the mapped sequence data as images, and treating variant calling as a special kind of image classification. It is reported that DeepVariant can outperform GATK in SNP calling. This demonstrates the potentials of deep learning in the sequence data processing domain. The DeepVariant approach raises a natural research question: can deep learning be applied to call other types of genetic variations from sequence data that are more complex than SNPs and short indels? In this paper, we provide a positive answer for this question: we show that deep learning can be used for accurately calling structural variations from sequence data.

Structural variation refers to relatively long genomic variation, such as deletion, insertion and inversion. Structural variation will lead to complications of many diseases [4], and many cancers are associated with genetic variation [5]. To be specific, we focus on calling long deletions (longer than 50 bp) in this paper. For deletion calling, there exists many approaches including Pindel [6], BreakDancer [7], DELLY [8], CNVnator [9], Breakseq2 [10], Lumpy [11], GenomeStrip2 [12], and SVseq2 [13], among others. Most of these approaches rely on one or multiple information (called signatures) extracted from mapped sequence data: (i) read depth, (ii) discordant read pairs and (iii) split reads. We note that there are also methods performing sequence assembly for deletion calling. While many of the existing methods have been used in large genomics projects such as the 1000 Genomes Project [14], there exists no single method that clearly outperforms other approaches.

In this paper, we present DeepSV, a deep learning based method for long deletion calling from sequence data. DeepSV builds on the general approach of DeepVariant by visualizing mapped sequence reads as images. The key technical aspects of DeepSV are the novel visualization techniques for CNN-based deletion calling and how to work with noisy training data. The visualization procedure combines all major aspects of features with regard to deletions: read depth, split read and discordant pairs. This avoids manual selection of features for classification. We demonstrate that DeepSV outperforms existing methods in calling long deletions on real sequence data from the 1000 Genomes Project. Our work extends the findings of DeepVariant by showing that deep learning can be useful for calling structural variations that are more complex than SNPs and short indels.

The rest of the paper is organized as follows. In Section 2, we survey the existing approaches for calling structural variations from sequencing data, and the application of the machine learning in the this subject. In Section 3, we present our deep leanring based SV calling method. In Section 4, we present the research results. In the last section, we provide discussions on the DeepSV approach.

## 2 State of the Art

Genomic deletions affect several aspects (called signatures) of the sequence reads mapped onto the given reference genome near the deletion site. (i) Read depth. Mapped read depth within a deletion is likely to be lower than those at wild-type sites. If the deletion is homozygous and the reads are mapped correctly (e.g. not mis-mapped due to repeats), read depth within the deletion should be close to zero. If the deletion is heterozygous, read depth within the deletion should still be lower than expected. Thus, low read depth is a signature of deletions. (ii) Discordant read pair. Consider paired-end reads that are mapped near the deletion with two ends being to the different sides of a deletion. Such read pair is called encompassing pair for the deletion. The mapped insert size (i.e. the outer distance of the two mapped reads of the pair) of an encompassing pair appears to be longer than usual due to the presence of the deletion. This is because the mapped insert size includes the length of the deletion on the reference genome. We say an encompassing read pair is discordant if the difference between its mapped insert size and the known library insert size is at least three times of the standard deviation of the library insert size. Otherwise, we say the read pair is concordant. The longer the deletion is, the more likely an encompassing pair becomes discordant. (iii) Split reads. When a read overlaps the breakpoints of a deletion, the read consists of two parts that are not contiguous on the reference: the part proceeding the left breakpoint and part following the right breakpoint. Such a read is called split read. Here, breakpoint refers to the boundary of the deletion on the reference genome. When a split read is mapped, the read cannot be mapped as a whole. Instead, it is mapped onto two discontinuous regions of the reference. These signatures reveal different aspects of structural variations. A main advantage of using split reads is that split reads can potentially reveal the exact breakpoints of the deletion. In contrast, read depth and discordant pairs cannot lead to exact breakpoints. See Figure 1 for an illustration.

**Fig. 1.**
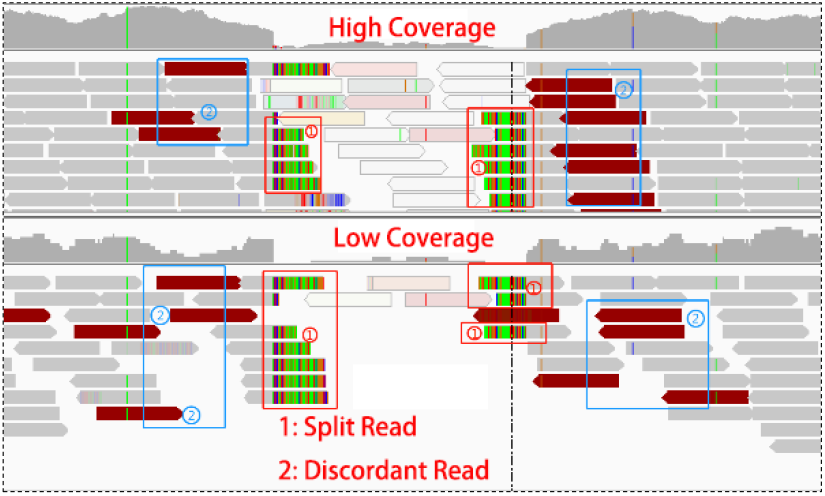
Above the image is high coverage data, and below is low coverage. The split reads are marked with ① and the discordant reads are marked with ②.

Most existing deletion calling methods utilize the above three types of signatures. Earlier methods often use a single signature. For example, BreakDancer uses only discordant read pairs. The original Pindel only relies on split-reads. A number of methods combine multiple signatures and obtained more accurate results. For example, several methods such as DELLY, MATE-CLEVER [15], and SVseq2 combined discordant read pairs and split reads. A main advantage of integrating multiple signatures is better utilizing the information contained in the sequence data. However, it is unclear what is the best way for integrating different signatures for deletion calling. A simple approach is weighting different signatures (e.g. a split read is weighted as 1 and a discordant pair is weighted as 2). Obviously, this leads to the issue of choosing the weights, which is often difficult in practice [16]. Indeed, it is known that parameter settings can affect the results of many existing SV callers [17].

A useful observation is that SV calling can be viewed as a classification problem. That is, for a candidate SV, we want to classify this candidate site to be either a true deletion (denoted as 1) or a false positive (denoted as 0) based on the given sequence data near the candidate site. Classification is an important subject of machine learning and there are many existing machine learning methods for classification. Usually classification involves two steps. First, a model is trained from training data. Second, the trained model is used to classify the test data. A main advantage of using a classification model is that there is no need to manually choosing the parameters; parameters are obtained from the training data. There are existing machine learning based approaches for SV calling, including GINDEL [18] and Concod [19]. While these machine learning based methods show promises in accurate calling of SVs, there are also difficulties faced by traditional classification methods. One of the most important issues for traditional classification is feature selection. That is, we need to determine what specific quantities to extract from sequence data to be used in classification. Due to the complex nature of structural variations, it is often unclear what are the best features.

Recently, deep learning is becoming increasingly popular. Deep learning approaches (such as convolutional neural network or CNN) have been applied to several important problems (e.g. image processing, computer vision, natural language processing, to name a few) and led to significant improvements in performance over existing methods. A main advantage of deep learning is that it reduces the need of feature engineering and can potentially better utilize the data. On the other hand, a main disadvantage is that deep learning usually needs more complex models that are more difficult to train than those in traditional approaches. Application of deep learning in sequence data analysis is in its infancy. A pioneering work in this area is Google’s DeepVariant. DeepVariant proposes a novel approach for sequence data analysis for the purpose of genetic variant calling: treating mapped sequence data as images and converting genetic variant calling to an image classification problem. As shown in Figure 2, mapped sequence reads have a natural visualization. Mapped sequence reads near a variant (say a short deletion) or wildtype site can be easily recorded as an image.

**Fig. 2.**
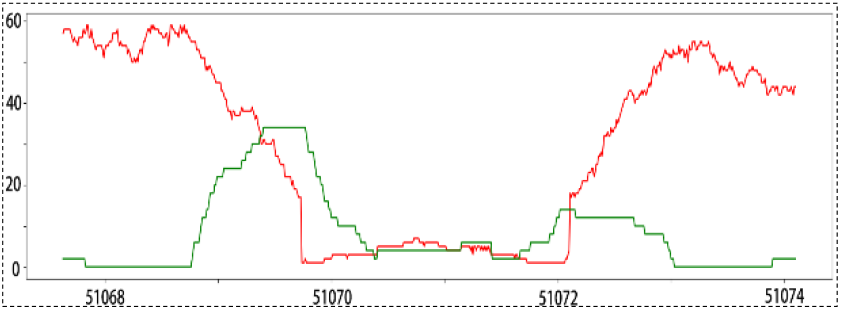
The red line is the read-depth value, the green line is split-read value and the insert size spaning the deletion region is larger than library value.

This image has different visual appearance at a variant from that from a wildtype. At a short deletion site, the image tends to have a gap. Moreover, read depth tends to be lower than that of the wildtype because short deletion may cause some reads unaligned. DeepVariant relies on the visual difference of images to perform classification for genetic variant calling. The DeepVariant approach leads to a natural question: can we use deep learning to call more complex genetic variants such as long deletions? SV calling can be somewhat more difficult than SNP or short indel calling. First, SNP calling is more localized: reads relevant to a SNP can be easily fit into a single image. Reads that are relevant for a long deletion can spread out. For example, two ends of a discordant read pair over a long deletion can be mapped to positions that are more than thousands of bases apart. Second, there are more signatures for long deletions than those for SNPs or short indels. For example, discordant read pairs are not associated with SNPs but are important for long deletions. Integrating these diverse set of signatures in visualization needs to be worked out. In this paper, we present DeepSV, a deep learning based method for calling long deletions from sequence reads, which addresses these difficulties.

## 3 Method

### 3.1 General Description of DeepSV

DeepSV is a deep learning based structural variation calling method. It is based on a new sequence reads visualization approach, which converts mapped sequence reads to images. DeepSV follows the general approach of DeepVariant. Different from DeepVariant, DeepSV aims to calling SVs (especially long deletions that are longer than 50 bp). There are two components in DeepSV: training and variant calling. Both components take mapped reads and the reference genome as input. For model training, DeepSV trains a convolutional neural network (CNN) model from sequence reads with known deletions. Similar to DeepVariant, CNN model training is based on visualizing mapped sequence reads near known deletion sites or at wildtype sites. The key technical aspect of DeepSV is how to train the CNN from sequence images near a deletion. Recall that long deletions have more complex signatures than SNPs or short indels. Note that deletion can be long and different regions of a deletion can be quite different in the visualized reads. For example, near the breakpoint, there is likely a sharp transition from high read depth to low read depth. In the middle of a deletion, there may be no such transition but the read depth can be lower than that near the breakpoint (See Supplementary Figure S1).

In order to accurately call deletions with precise breakpoints, it is important to separate these cases. Moreover, to accommodate various signatures of a deletion, DeepSV implements a visualization procedure that takes advantage of the rich information contained in an image to integrate various signals. In a typical color map, there are 8 bits for red, green and blue and so there can be 256 choices for each color. Therefore, one can use various combinations of the three basic colors to represent the configuration of the mapped reads. For example, a pixel corresponding a base of a mapped read can be affected by multiple factors such as whether the read is split read, the quality of the read, whether there is a discordant read pair and so on. When the CNN model is trained, the model is used to call deletions from the sequence images.

### 3.2 DeepSV workflow

Figure 3 shows the overall workflow of DeepSV. DeepSV is composed of three parts. In the first part (Fig.3.a), DeepSV begins by finding candidate deletions in reads aligned to the reference genome using clustering. In the second part (Fig.3.b), the deep learning model is trained using a pileup image of the reference and reads around each candidate variant. In the third part (Fig.3.c), the model is used to call the variants.

**Fig. 3.**
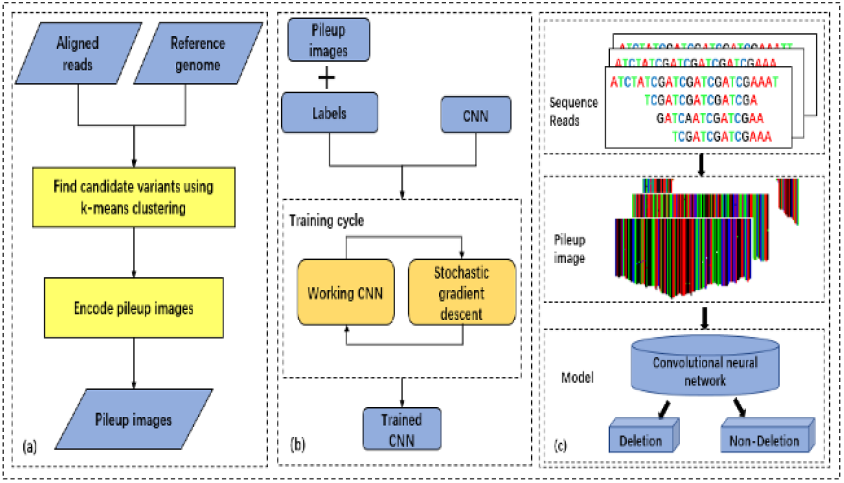
Overview of the DeepSV. DeepSV extracts candidates training set from the original sequence, and then uses a gradient descent algorithm to train CNN network, and finally generates images according to gene sequence for calling deletions. The gradient descent algorithm is an optimization algorithm commonly used in machine learning and artificial intelligence to recursively approximate the minimum deviation model. (a) Finding candidate variants and Encoding pileup images. (b) Training CNN model. (c) Calling deletions.

### 3.3 Model training

To train a classification model, DeepSV takes the aligned sequence reads in BAM (binary sequence alignment/map format) file and a VCF (Variant Call Format) file which contains the known deletions. We partition the reference genome into consecutive non-overlapping windows of 50bp. The aligned reads in the pileup format are converted into an image.

Model training needs a set of labeled training samples. For image-based deletion calling, we need two sets of images: images from the deletion regions (labeled as 1) and images from the wild-type regions (labeled as 0). In principle, since the deletions are given in the VCF file, creating training sample is straightforward. Mapped reads have the natural pileup form and can be easily converted to images. 1-labeled images are taken within the known deletions and 0-labeled images are from outside the deletions. Once the training data is obtained, one may train a CNN. In practice, however, there are three challenges for developing a deep learning based deletion caller.

- Currently known deletions from large genomics project (e.g. the 1000 Genomes Project) are usually noisy. While many known deletions have relatively accurate breakpoints, some deletions in the VCF have only a single breakpoint. Also some given breakpoints don’t match well the aligned reads when one visualizes the reads in the IGV tool [20]. Noise in the known deletions in the training data can significantly affect the performance of DeepSV.
- Images should be formed to improve the classification performance. The images need to integrate multiple sources of information (i.e. read depth, discordant pairs and split read). The images need to carry strong information on whether an image is from a deletion or not. Our experience shows that this isn’t a trivial task: in our initial attempt, an alternative visualization approach performs poorly for deletion classification.
- CNN model selection is important. It is well known that choosing the right CNN model has a large impact on the classification performance. Choosing CNN models often involves trial and error.

#### 3.3.1 Image labeling on noisy data

In order to obtain reliably labeled training images, we first preprocess the sequence reads and the given deletions. The basic idea is filtering out deletions that don’t match the sequence reads well. The noise filtering is performed based on a k-means clustering procedure. Consider a given deletion with known breakpoints. The clustering treats each genomic position near or within this deletion as a point in three-dimensional space. Here, the three axes are read depth, discordant read pair count and split-read count at a specific position. Intuitively, signature values of genomic positions inside a deletion tend to be similar (e.g. have smaller read depth) and thus these positions tend to form a cluster. Similarly, positions from the upstream regions of the deletions tend to form a cluster, and so do the positions from the downstream regions of the deletions. See Figure 4 for an illustration. Therefore, we let k=3 and run k-means clustering to cluster the positions into three categories. The three clusters are denoted as *S*_1_, *S*_2_, *S*_3_ which correspond to the upstream, deletion and downstream regions respectively.

**Fig. 4.**
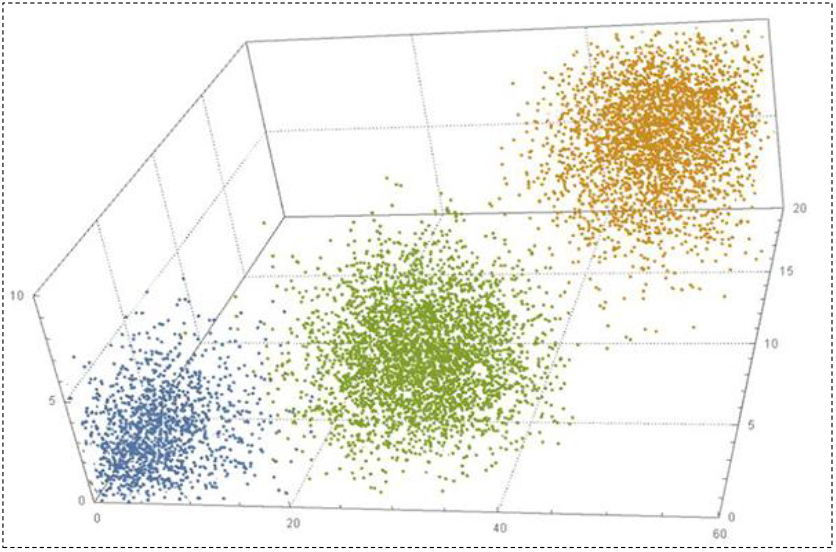
x aixs: read depth value, y axis: split read value, z axis: discordant value. From the figure, The three values (x, y, z) of the deletion region are relatively small and close to the orgin(blue points). The number of split reads (y value) and discordant reads (z value) are more near the breakpoint(green points). The read depth value (x value) are higher on non-deletion region(yellow points).

The clustered positions are used to eliminate false positives, find exact breakpoints and generate training labels as follows. Note that the lower read depth is, the higher split read count and discordant read count are for deletions. And upstream and downstream regions tend to have higher depth and low split read counts and less discordant read count than deletion regions. So we compute a feature value *m* for each position, where *m* is equal to the read depth minus split read count and discordant read count. We compute the average feature value of all positions in each of the three clusters *S*_1_, *S*_2_, *S*_3_, which we denote as 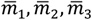. We let 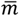 be the minimum of the tree means. If the positions from the cluster with the minimum mean are largely between the points of the other clusters, then this deletion is considered to be a true deletion. Otherwise, it is a false positive. Our experience indicates that this removes many false positive.

Sometimes the given breakpoints are not very accurate. This can lead to wrong labeling of the training images, especially near the boundary of the deletion. To find exact breakpoints, we consider the minimum mean cluster *S*_2_ with mean 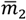. DeepSV sorts the positions of *S*_2_. The minimum and maximum positions, denoted as *β*_1_ *and β*_2_, are treated as initial breakpoints. There are three cases for the interval [*β*_1_, *β*_2_].

i. [*β*_1_, *β*_2_] is close to the given breakpoints.
ii. [*β*_1_, *β*_2_] doesn’t include the given breakpoints due to the length of the deletion being too long.
iii. [*β*_1_, *β*_2_] doesn’t match the given breakpoints at all. We set two pointers *p*_1_ *and p*_2_ to *β*_1_ *and β*_2_ respectively. If the feature value of *p*_1_ is less than 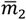, then *p*_1_ is moved to the left by one. Similarly, if the feature value of *p*_2_ is less than 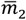, then *p*_2_ is moved to the right bye one. When this process finishes, the two pointers provide the estimate of the two breakpoints of this deletion.

##### Dealing with noise

Real sequence data tends to have significant noise, which can make the clustering perform poorly. The following lists several such cases.

i. The read depths fluctuate and some positions have read depths that are either too high or too low than expected. For example, read depths at some sites in the non-deletion region can be very low, while read depths of some sites in a deletion can be very high. Read depth near the breakpoints fluctuates significantly.
ii. The difference on discordant paired-end reads between a deletion and a non-deletion is not obvious when coverage is low.
iii. Split read counts and discordant read counts are inversely proportional to the depth value near the deletion breakpoints. In the clustering algorithm, the data clustering for the same trend tend to perform well. Data with different trends in different aspects tend to make clustering work poorly.

To address the above three issues, DeepSV uses the following techniques to reduce the effect of noises. First, DeepSV uses a 61 bp long sliding window to filter the read depths. DeepSV uses the following filter formula: 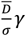. We let the breakpoints of a deletion to be [b1, b2]. The mapping read depths within the window are *d*_1_, *d*_2_, …, *d*_61_(computed by SAMtools [21]), and the 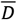 is the average of the read depths of the window. σ is the standard deviation of *d*_*i*_ and γ represents a coefficient. 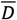 can indicate the situation where the read depth is high in the non-deletion region or low in the deletion region. σ reflects the fluctuations of depth. The value of γ is chosen to amplify the trend of the depth values in the window. Our experience indicates that this filtering step reduces the effect of the noise in the data, and improves the performance of the clustering. Now we consider discordant reads and split reads. Since the split read count and the discordant read count can be inversely proportional to the read depth near the breakpoints, we use the negation of the discordant read counts and split read counts (instead of the reads counts themselves). This is to ensure that each feature used in the clustering has the same trend for deletions or non-deletions. This improves the performance of the clustering. In order to ensure that the boundary of clustering is as close as possible to the breakpoints and improve the accuracy of called deletions, DeepSV uses a modified Euclidean distance formula for the distance measure between two points used in the clustering. Euclidean formula can better reflect the distance relationship of space vectors. It is a commonly used distance formula in clustering algorithm. Considering that each coordinate in multidimensional vector contributes equally to Euclidean distance and they often have random fluctuations of different sizes, we have improved the formula by adding weight adjustment.

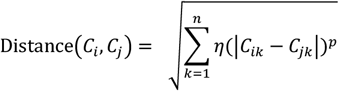

*C*_*i*_ *and C*_*j*_ represent two points of cluster set. *C*_*ik*_ *and C*_*jk*_ are the components of vectors in the form of (sites, normalized read depth, negated split read count, negated discordant read count). When the *C*_*ik*_, *C*_*jk*_ values are the normalized depth depths, η is a fixed number that is greater than one. For discordant read count and split read count, η is set to one. The constant p is a fixed even number.

#### 3.3.2 Visualizing mapped sequence reads

So far, we have labeled regions along the reference genome to be either deletion or non-deletion. We now describe how to create images for each region. This is a critical step because real sequence data tends to be noisy. If the visualization approach is not chosen properly, the CNN may not capture the underlying information about the deletion from the created images.

Recall that an image is composed of pixels, and each pixel has (R, G, B) three-primary colors. DeepSV takes the following simple approach for visualizing the reads: each nucleotide (i.e. A, T, C, or G respectively) is assigned one of these base colors: red (255,0,0), green (0,255,0), blue (0,0,255) and black (0,0,0) respectively; then the base color is slightly modified to integrate the various signatures on deletions. DeepSV considers all the aligned bases (i.e. a column in the image) at a position of the current region. Since most columns have the same base, the bases from one column tend to have similar colors. This gives the visual appearance of column-based images. See Figure 5 for an illustration. Our experience shows that such column-based images reveal the key aspects of deletions.

**Fig. 5.**
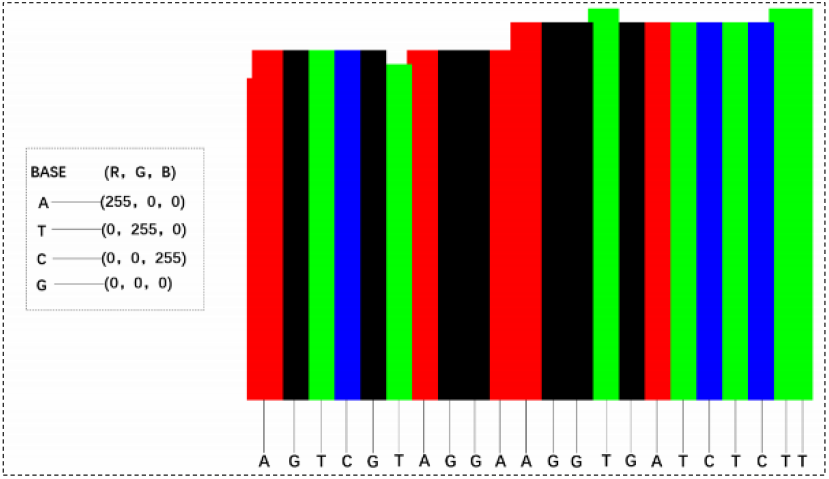
Each base is assigned one base color, and each color is composed of three primary colors. According to the characteristics of alignment, the bases at each site are assigned different colors.

##### Integration of deletion signatures

Recall that there are various signatures on deletions (i.e. read depth, discordant pair and split read). Read depth is naturally represented by the pile-up images. DeepSV integrates the other two types of signatures by slightly modifying the base colors of the mapped bases based on the signatures. Note that such modification is usually mild and does not destroy the column-based appearance of images. Each read contains multiple aspect of information, e.g., whether it belongs to discordant paired-end reads and whether it is split read. DeepSV uses the following combination of features to determine the color of each mapped base. More specifically, the color of a mapped base is determined by the four quantities, which describe the discordant read pair and split read information at the position. These quantities are explained in Table 1. The sum of these four quantities provide the auxiliary components of the coloring. To see how these four quantities are used to decide the color of a mapped base, we consider the following example. Consider a column of aligned bases (which are say all A’s). We first count the number of discordant paired reads (which is say 10), and the number of split reads (which is say 0) that overlap this column. This leads to a color setting (255,10,10) for all bases in the column. We then consider each base of the column one by one. For this, we find the four features (is paired, concordant/discordant, mapping quality and map type). Say these features have values (1,1,1,0). This leads to a binary number 13=10+1+1+1+0.

**Table 1.** Visualizing an aligned base. Feature: information carried by the base on deletions.

|  | Feature | Description |
| --- | --- | --- |
| 1 | is-paired/is-not-paired | Is this read paired (1) or single (0)? |
| 2 | concordant/discordant paired | Is this read discordant (value 1) or not (value 0) |
| 3 | mapping-quality | Is the mapping quality higher than 20? Value is 0 or 1. |
| 4 | map-type | Is this read split (value 1) or not (value 0)? |

As shown in Table 1, each of the four bases has four binary feature values. This gives total 64 combinations. We show an example (See Supplementary Table S1) that describes 64 value combinations and color scope. The color range of A base is (255, 0, 0) ∼ (255, 235, 235), T base is (0, 255, 0) ∼ (235,255, 235), C base is (0, 0, 255) ∼ (235, 235, 255), and the G base is (0, 0, 0) ∼ (235, 235, 235).

Because the range of deletion length is from 50bp to more than 10kbp, a single pileup image cannot cover an entire deletion and discordant pairs may not be contained in a single image. DeepSV divides the deletion region into equal length regions. DeepSV generates the fixed length images for to each region. This is illustrated in Figure 6.

**Fig. 6.**
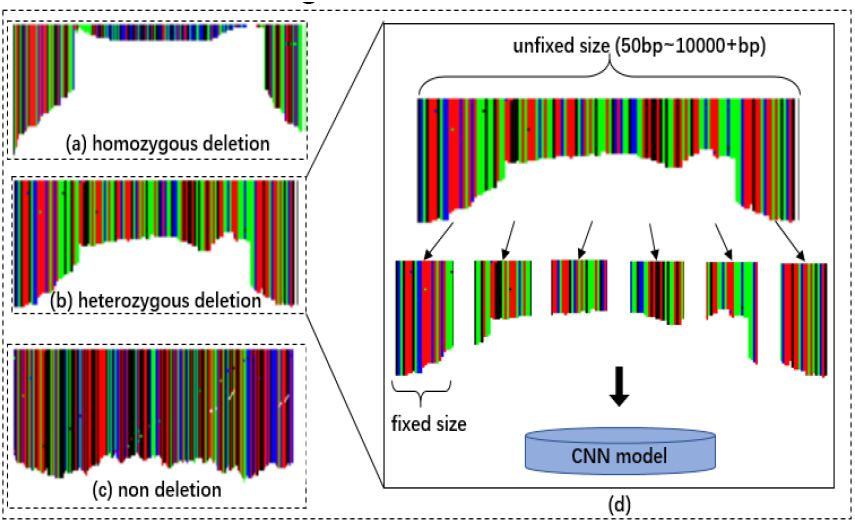
These images contain different features. Each vertical colorful bar represents bases aligned to this site. The length of the vertical bar becomes lower in the deletion region but higher in the non-deletion region.

#### 3.3.3 Training and Validating Model

DeepSV trains the CNN model with real sequence reads and benchmarked deletions. Tensorflow [22] is used to construct the convolutional neural network. All genome data analysis is performed on a Linux server with 1080Ti GPUs [23] and a platform of Digits [24]. The parameters setting of model on deletion calling are given in the supplemental materials (section 3).

##### Picture labeling and normalization

The nucleotide sequence of the entire deletion region is divided into 50bp regions. DeepSV generates images of 256 by 256 for these 50bp and labels each image as 1. Similarly, a non-deletion region is divided into 50bp regions to generate images with labels of 0. We use all the labeled images in CNN training and use the trained model to call the deletion of the test data. The labeled images are shown in Supplementary Figure S6. The deletion calling process is shown in Supplementary Figure S7.

## 4 Result

We now validate the performance of DeepSV using real data from the 1000 Genomes Project. The called deletions released by the 1000 Genomes Project (phase three) are used as the ground truth for benchmark. The data we use in this paper consist of 40 BAM (binary sequence alignment/map format) files with 20 individuals on chromosomes 1∼22. The average insert length is 456bp, and the standard deviation is between 57bp∼78bp. The average coverage is 10X and 60X. These individuals from YRI, CHB and CEU (representing three different populations). DeepSV needs training data. We use about half of data for the purpose of training, and use the remaining data for testing. The training data set and the test data set are divided according to the following two criteria: (i) ensuring that the training data is sufficient for the model to converge. (ii) ensuring that the test data is sufficient to cover various targets to be detected.

Under the premise of satisfying the above two criteria, the ratio of the training set and the test set can be adjusted according to the actual situation. In this experiment, the training set and the test set are each 50%. For training, we use the data from chromosomes 1 to 11 of these 20 individuals. For testing, we use the data from the chromosomes 12 to 22. Data used in the experiments is given in the Supplementary (Table S3).

We compare DeepSV with other eight tools including Pindel, BreakDancer, Delly, CNVnator, Breakseq2, Lumpy, GenomeStrip2, and SVseq2. To show the advantage of deep learning, we also compare with an existing machine learning based method, Concod. We examine various aspects of deletion calling by DeepSV and other tools, including the accuracy of calling deletions of different sizes, breakpoint accuracy, impact of sequence data coverage, and the effect of model’s activation on precision and loss. Due to the space limit, some results are given in the supplemental materials.

### 4.1 Calling deletions of different sizes

We first evaluate the performance of DeepSV for calling deletions of various sizes. We use the deletions on chromosomes 12 to 22 from 20 individuals from the 1000 Genomes Project as the benchmark. Figure 7 shows the deletion distribution of these benchmarked deletions. We divide the lengths of deletions into five categories. The deletion length roughly follows a normal distribution.

**Fig. 7.**
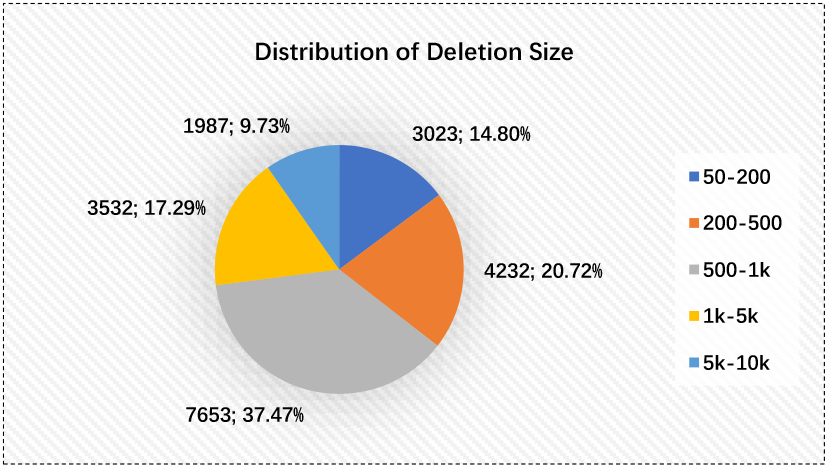
Deletion length distribution for deletions in the chromosomes 12 to 22. The deletion length obeys the normal distribution

We compare the accuracy of deletion calling of different deletion sizes on DeepSV and other calling tools with low or high coverage data. To measure the performance of deletion calling, we use the following statistics: precision (P), sensitivity (S), and the F-score. The results on low coverage are shown in Supplementary Table S4. The results on high coverage are shown in Supplementary Table S5.

#### 4.1.1 DeepSV for long deletions: the effect of complex SV

We now take a closer look at the performance of DeepSV on calling long deletions. Deletion can be classified into homozygous and heterozygous deletion. Moreover, there can exist other types of structural variations (e.g. insertion, translocation and inversion) near the deletion region. This type of structural variation is called complex SV. The importance of complex SV has been recently noticed in the literature [25]. A deletion of longer size is more likely to be a complex SV than shorter deletions. When a deletion is complex, the images created by DeepSV tends to be less clear cut than those from a simple deletion. See Figure 8(c) for an illustration. To evaluate the performance of DeepSV on calling different types of deletions, we study how DeepSV’s performance changes when the number of other types of structural variations (e.g. insertions, translocations and inversions) increases. The results are shown in Figure 8(d). Our results show that the accuracy of calling homozygous deletions is not affected by the presence of other types of SVs in general. The same roughly holds for heterozygous deletions. For complex deletions, the presence of other SVs reduces the calling accuracy significantly.

**Fig. 8.**
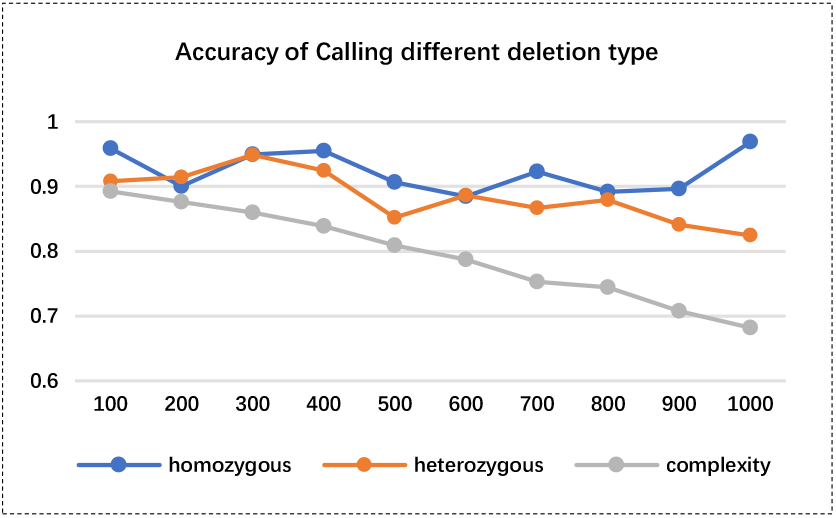
Accuracy of calling deletions for different type. Horizontal axis: number of other types of SVs (e.g. insertion/translocation/inversion) near the deletions. Vertical axis: deletion calling accuracy (part d).

### 4.2 Breakpoint accuracy

In this section, we compare the performance of DeepSV with other tools on breakpoint accuracy. Figure 9 shows the distance between the detected and true breakpoints on the all genome data. Once again, breakpoint predictions given by DeepSV are closest to the true breakpoint positions. In many cases, the predicted breakpoint positions of DeepSV are only up to a few base pairs away from the true breakpoint positions. Predictions from other tools are usually further away from the true breakpoint positions. Among them, SVseq2, CNVnator, Lumpy, and BreakSeq2 appear to have the lower breakpoint accuracy than other tools. GenomeStrip2 has the highest resolution in the tools we compared. Our results suggest that DeepSV appears to capture the key characteristics of breakpoints from the images.

**Fig. 9.**
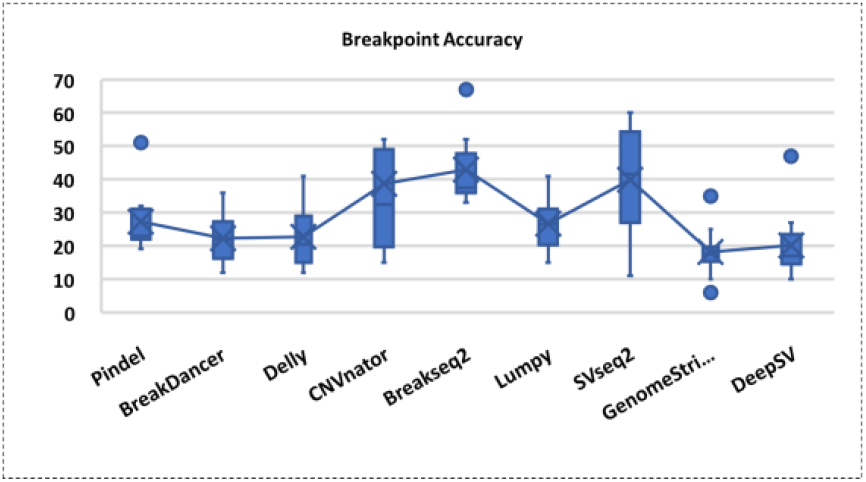
Breakpoint accuracy on different detecting tools. The y-axis represents the distance between detected breakpoints and true breakpoints.

### 4.3 Impact of reads coverage

Sequence reads coverage can have a large effect on deletion calling. Reads coverage also affects the performance of DeepSV in both training and testing. In order to evaluate the performance of DeepSV on datasets with different coverage, we perform down sampling for the original high coverage data. SAMtools with option “view -s” is used for down sampling, and four datasets are generated with average coverage 48×, 36×, 24×, 12× respectively. Table 2 shows the precision, sensitivity of DeepSV and the other eight tools on the genome data. In the table, we can see DeepSV has the highest precision compared with all other tools at various coverage. The sensitivity of DeepSV is overall comparable with the best of the other tools.

**Table 2.** Precision, sensitivity of multiple tools on different coverage data. P indicates precision. S indicates sensitivity.

| Tool | 12 × |  | 24 × |  | 36 × |  | 48 × |  |
| --- | --- | --- | --- | --- | --- | --- | --- | --- |
|  | P | S | P | S | P | S | P | S |
| Pindel | 0.18 | 0.80 | 0.45 | 0.50 | 0.33 | 0.60 | 0.40 | 0.48 |
| BreakDancer | 0.50 | 0.86 | 0.39 | 0.51 | 0.68 | 0.63 | 0.70 | 0.46 |
| Delly | 0.30 | 0.83 | 0.43 | 0.57 | 0.69 | 0.51 | 0.76 | 0.38 |
| CNVnator | 0.17 | 0.18 | 0.36 | 0.20 | 0.61 | 0.59 | 0.75 | 0.74 |
| Breakseq2 | 0.69 | 0.71 | 0.71 | 0.75 | 0.80 | 0.79 | 0.86 | 0.85 |
| Lumpy | 0.13 | 0.87 | 0.21 | 0.93 | 0.32 | 0.82 | 0.42 | 0.73 |
| GenomeStrip2 | 0.36 | 0.42 | 0.82 | 0.69 | 0.86 | 0.78 | 0.75 | 0.60 |
| SVseq2 | 0.57 | 0.28 | 0.43 | 0.22 | 0.30 | 0.33 | 0.23 | 0.36 |
| DeepSV | 0.72 | 0.81 | 0.85 | 0.89 | 0.91 | 0.92 | 0.93 | 0.90 |

### 4.4 Deletion calling for various frequencies

To see whether DeepSV performs consistently on different population deletion frequencies, we now show the results of DeepSV on various deletion frequencies within the population and compare with the other eight tools. Note that, the precision cannot be calculated because the deletion frequency for the false positive deletions called out by each tool is unknown. Therefore, we only list the sensitivity here. Supplementary Table S6 shows the performance of each tool on different deletion frequencies. The results show that DeepSV outperforms the other eight tools for different deletion frequencies. The sensitivity of most tools increases as deletion frequencies increase. GenomeStrp2 has the better sensitivity for medium deletion frequency of 6-10. CNVnator has the lowest sensitivity for medium deletion frequency of 1-5.

### 4.5 Deletion calling for an individual not in training

So far, we use data from 20 individuals where half chromosomes are used for training and the other half for testing. Since training and testing are on different chromosomes, testing is considered to be independent from training. To further validate our method, we now show deletion calling performance for an individual (NA12891) that is not used in the training. The results for this individual with various coverage are shown in Figure 10. We can see that DeepSV performs well for this new individual with performance similar with those from other individuals.

**Fig. 10.**
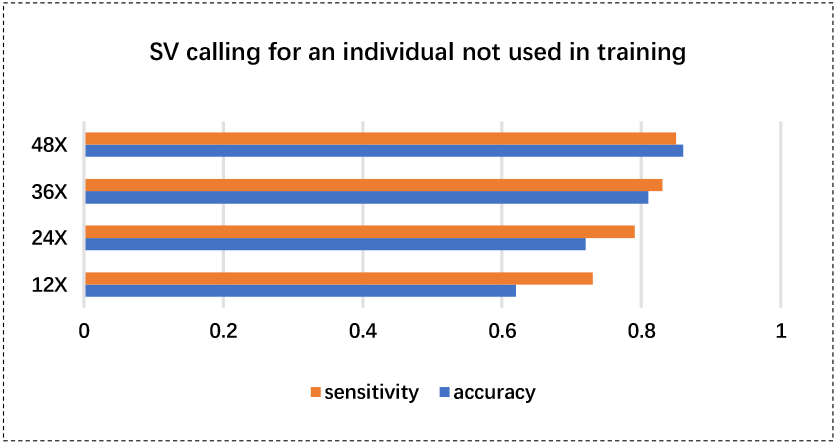
Performance of deletion calling for NA12891. To verify the DeepSV’s validity, we test the individual of NA12891 that is independent from traning and testing.

### 4.6 Machine learning and deletion calling

There are existing methods that use other types of machine learning approaches for deletion calling. Concod is one such example. Concod is based on manual feature selection. It performs the consensus-based calling with a support vector machine (SVM) model. We now compare DeepSV with Concod in deletion calling. Both methods need model training. We compare the model training accuracy and loss, as well as the running time of the two methods. Here, model training loss is the model misclassification error on the training data on a trained model. That is, we first use training data to train the model. Then we treat the training data as the test data to see if the model classifies the training data correctly. Note that there is an overfitting issue: a model classifies training data well may not generalize to test data. Nonetheless, a good machine learning model should have small training loss. The results are shown in Figure 11(a,b).

**Fig. 11.**
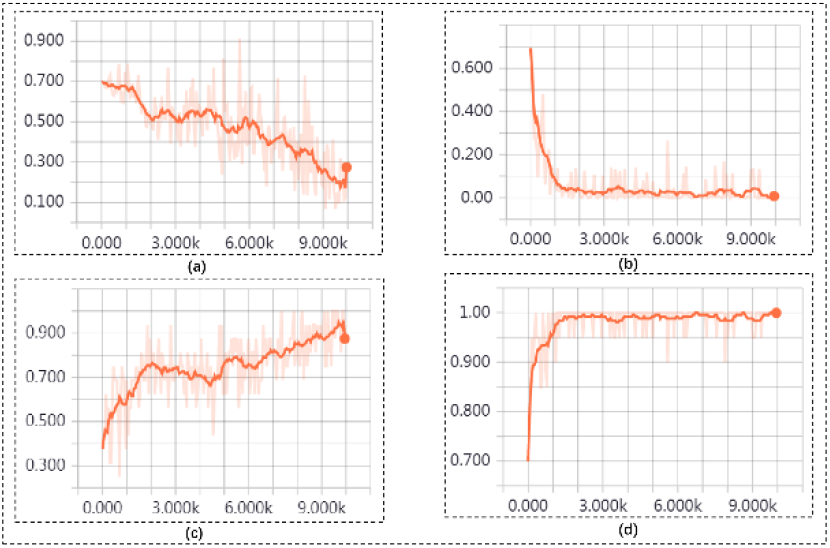
Compared to traditional machine learning method (e.g., Concod), DeepSV shows a more stable state. The training loss of Concord(a) and DeepSV(b). The training accuracy of Concod(c) and DeepSV(d).

As shown in Figure 11(c,d), DeepSV outperforms Concod in training accuracy. This suggests that DeepSV is better in deletion calling than Concod. For running time, Concod has smaller training time when the number of training samples is small. When the number of training samples is large, Concod takes longer time than DeepSV in training. The results are shown in Figure 12.

**Fig. 12.**
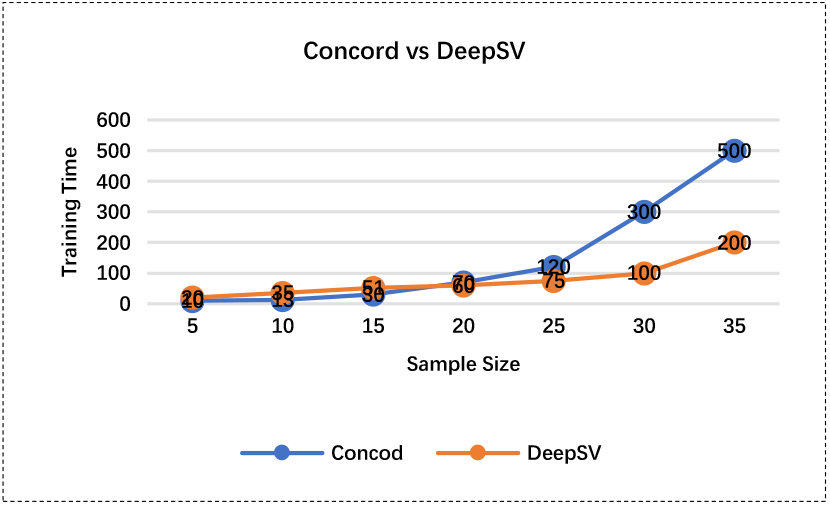
The relationship between training time and sample size about Concod and DeepSV.

## 5 Discussion

On the high level, calling genetic variations from sequence reads is a process of extracting information in the reads that are relevant to the variations. Google’s DeepVariant approach shows that such information can be effectively extracted from color images constructed from the reads. Amain advantage for this visualization based approach is that it offers an intuitive way to convert variation calling to image classification. This naturally leads to deep learning, which is currently the leading approach for image classification. Our DeepSV method extends this high-level approach to the case of structural variations. In particular, our results show that the visualization approach can be used for more complex genetic variations. Our results show that the overhead for visualizing sequence reads is low. The images contain useful information on structural variations. Deep learning can outperform other traditional machine learning since it doesn’t depend on manual selection of features. This may allow deep learning based methods to better utilize the data.

Usually deletion calling performance is affected by the type of data (e.g. sequence coverage and deletion lengths). A method can perform well for some type of data (say high coverage on short deletions) but doesn’t perform well for other types (say low coverage on long deletions). Our results show that DeepSV performs well in almost all the settings. This indicates that DeepSV integrates and effectively uses various sources of information in the sequence data in our simulation. Similar to other supervised machine learning methods, DeepSV needs labeled training data. The more accurate the training data is, the better DeepSV performs for deletion calling. With large-scale genomics projects such as the 1000 Genomes Project, high-quality training data is becoming available. Most existing methods for SV calling don’t use the known genetic variations. This may indicate a potential loss of information and missed chances. DeepSV offers a natural way for using these benchmarked SVs for accurate calling of novel SVs.

This paper focuses on deletion calling. We note that there are other types of structural variations such as long insertions, inversions and copy number variations. A natural research direction is developing methods for calling these types of SVs with deep learning. This will need more thoughts on methodologies. For example, long insertions are different from deletions in that the inserted sequences are not present in the reference genome. This makes the reads visualization more difficult since DeepSV currently visualizes reads on the reference. Such issues need to be resolved in order to apply deep learning to these types of SVs.

## Supporting information

All Supplemental Materials of Manuscript

## Declarations

## Acknowledgements

We would like to thank Kai Ye for useful discussions.

## Author’s contributions

Conceived and designed the experiments: YW JG LC. Performed the experiments: LC. Analyzed the data: LC. Contributed reagents/materials/analysis tools: YW LC. Wrote the paper: LC YW JG.

## Availability of data and materials

We use the real data from the phase three of the 1000 Genomes Project. The data are downloaded from: ftp://ftp.1000genomes.ebi.ac.uk/vol1/ftp/phase3/data/. The software and sample result as part of this project are readily available from GitHub at https://github.com/CSuperlei/DeepSV.

## Ethics approval and consent to participate

Not applicable.

## Consent for publication

Not applicable.

