## Supplementary material for "DeepSV: Accurate calling of genomic deletions from high-throughput sequencing data using deep convolutional neural network": All Supplemental Materials of Manuscript

### 1 High level approach

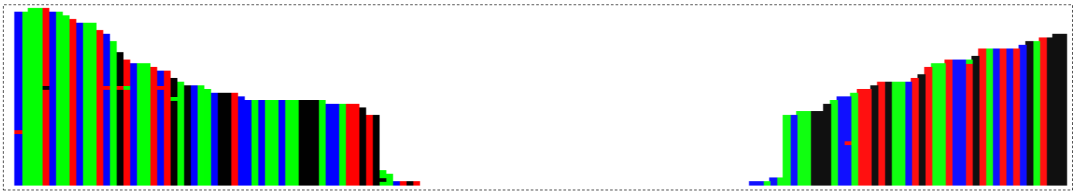

**Fig.S1.** The outline of the deletion region. The read depth is high on both sides, but is lower in the middle of a deletion. Each vertical bar represents an alignment site and different colors represent different bases. The color values merge the alignment features.

### 2 Feature combinations and color expression

Features are the key step for calling deletions. Good features contain information to distinguish true and false ones. The DeepSV combines 64 features which can effectively and comprehensively represent the properties of deletions. In these process, we not only consider the features that indicate the presence of deletion, we also collect features that imply the absence of deletions. Above all, DeepSV makes each feature to incorporate into the colors of R, G, B. In the Table S1, we exhibit these 64 features and describe the meaning of each characteristic and color.

**Table S1.** Feature combinations and corresponding Base's color. We list the interval values of colors, which represent the combination of different features.

| **Base** | **Feature Describe** |
|---|---|
| A<br>T<br>C<br>G | is-paried, concordant, mapping_quality>20, map_type=0 |
| | is-paried, concordant, mapping_quality>20, map_type=1 |
| | is-paried, concordant, mapping_quality<20, map_type=0 |
| | is-paried, concordant, mapping_quality<20, map_type=1 |
| | is-paried, discordant, mapping_quality>20, map_type=0 |
| | is-paried, discordant, mapping_quality>20, map_type=1 |
| | is-paried, discordant, mapping_quality<20, map_type=0 |
| | is-paried, discordant, mapping_quality<20, map_type=1 |
| | is-not-paried, concordant, mapping_quality>20, map_type=0 |
| | is-not-paried, concordant, mapping_quality>20, map_type=1 |
| | is-not-paried, concordant, mapping_quality<20, map_type=0 |
| | is-not-paried, concordant, mapping_quality<20, map_type=1 |
| | is-not-paried, discordant, mapping_quality>20, map_type=0 |
| | is-not-paried, discordant, mapping_quality>20, map_type=1 |
| | is-not-paried, discordant, mapping_quality<20, map_type=0 |
| | is-not-paried, discordant, mapping_quality<20, map_type=1 |
| **A** | 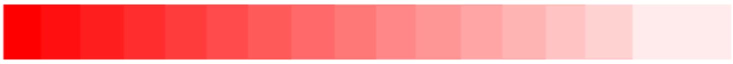 |
| **T** | 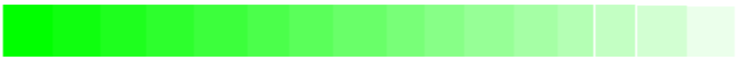 |
| **C** | 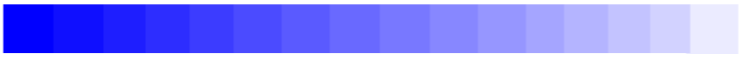 |
| **G** | 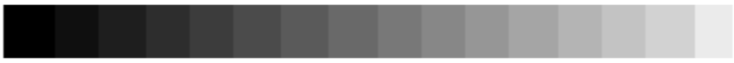 |

Different colors represent different features and each base has a main color. The lighter the base's color indicates, the worse the quality of base aligns.

### 3 The parameters setting of model on deletion calling

#### 3.1 Construction of Convolution Neural Network

The architecture of the convolutional neural network is shown as Table S2. The grid search and 10-cross validation are used to find the optimal parameters, and the default stochastic gradient descent algorithm is used. The network training parameters are set as following: the training epochs are set 30 and the learning rate is set 0.001.

**Table S2.** The CNN consists of 12 layers and each layer has its own function. Convolutional layer is responsible for learning picture features. The Pooling layer function is downsampling. Flattern layer plays an excessive role. The Fully connected layer is used as the final classification.

**Cov: convolutional layer. Pool: max pooling layer. Fc: fully connected network layer. Leaky ReLU: retified linear function.**

| Layer | Type | Activation | Feature Maps | Filter Size | Output Size |
|---|---|---|---|---|---|
| 0 | input | | | | 256*256 |
| 1 | Cov | Leaky ReLU | 96 | 11*11 | 246*246 |
| 2 | Pool | Max_Pool | 96 | 3*3 | 82*82 |
| 3 | Cov | Leaky ReLU | 256 | 5*5 | 78*78 |
| 4 | Pool | Max_Pool | 256 | 3*3 | 26*26 |
| 5 | Cov | Leaky ReLU | 384 | 3*3 | 24*24 |
| 6 | Cov | Leaky ReLU | 256 | 3*3 | 22*22 |
| 7 | Pool | Max_Pool | 256 | 3*3 | 8*8 |
| 8 | Flatten | | | | |
| 9 | Fc | Leaky ReLU | 512 | 512 | 1*512 |
| 10 | Drop_out | | | | |
| 11 | Fc | Leaky ReLU | 512 | 512 | 512*1 |
| 12 | Drop_out | | | | |
| 13 | Fc | | 2 | 2 | 1*2 |

#### 3.2 Impact of different DeepSV training parameters

We need to point that different layers and parameters' configuration will affect the model training effect. When we train the model for classification, the training parameters are adjustable according to the experience and the requirement for accuracy and loss. Figure S2 shows the results of model's loss rate with different parameters on training model. We try different layers associating with various parameters for training model. Each line in Figure S2 indicates the model effect with specific parameters of epoch/learning rate. The epoch means the total number of training times on the training set. The learning rate indicates the rate of weights update, which set too large will make the results exceed the optimal value but too small will make the loss of decline quite slow. From the chart, we can see that when the epoch is set 30 and learning rate is set 0.001, the loss is lowest. The user can set parameters according to the specified requirements of accuracy and loss.

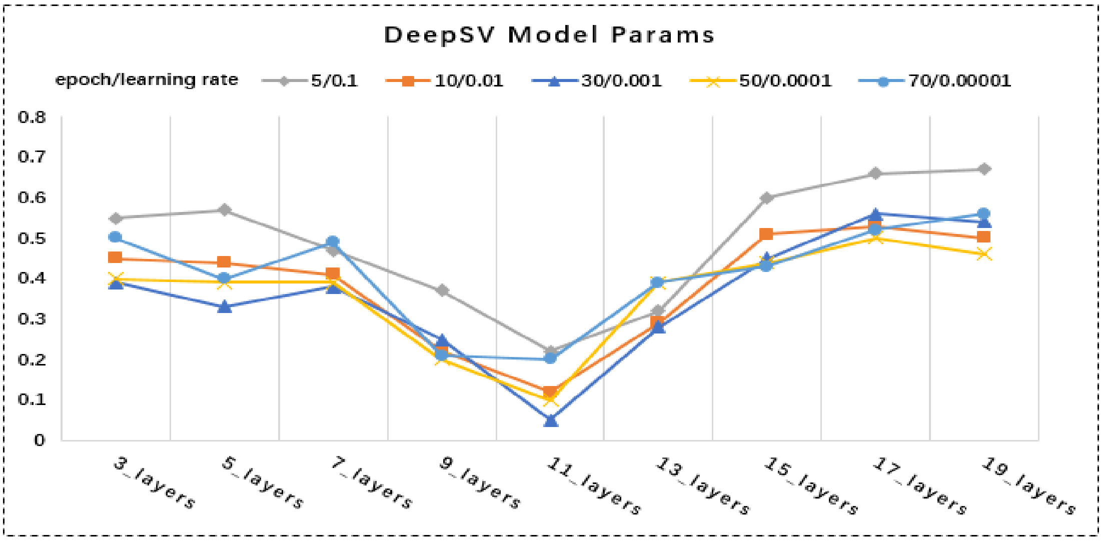


**Fig.S2.** The line diagram shows model convergence on different parameters. The x-axis indicates the number of network layers. The y-axis represents loss with different parameters of epoch/learning rate.

**3.3 Effect of model's settings on precision and loss**

When we train the CNN model, if the model is used to predict a certain image and it turns out that the activation matrix of a feature map in the convolutional layer looks basically the same as the original input, this is an indication that some problems occur because this feature map does not learn much useful information. The reason for this problem is the improper setting of the activation. Figure S3 shows the effects of different activation on model's accuracy and loss. From this figure, we can see that Leaky ReLu activation is set in the model with faster convergence rate and smaller error fluctuation on the final forecast. The Tanh and ReLU activation and loss will not influence final prediction but the convergence curve fluctuates more. The Sigmoid activation is not suitable to be used in CNN.

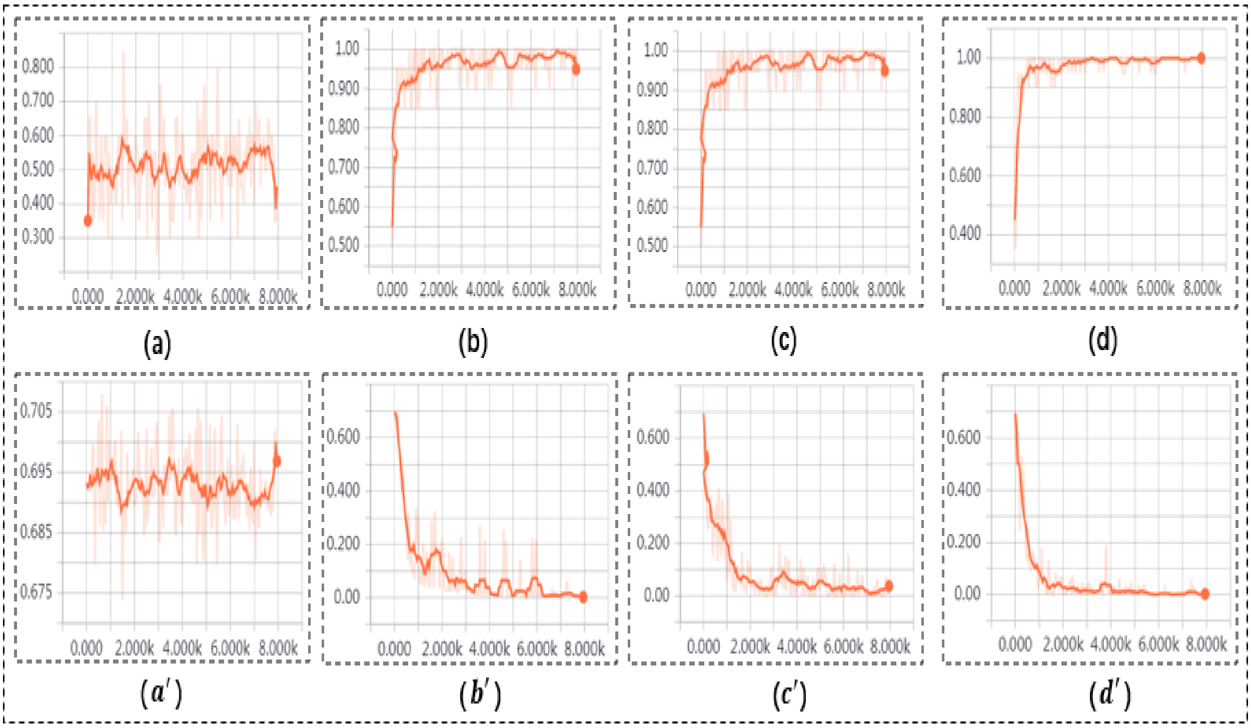


**Fig.S3.** The result of different activation with the convolution layer. The upper part of the picture represents training accuracy, and the lower part of the picture represents loss. Accuracy:(a) Sigmoid (b) Tanh (c) ReLU (d) Leaky ReLU. Loss:($a'$) Sigmoid ($b'$) Tanh ($c'$) ReLU ($d'$) Leaky ReLU

#### 3.4 Visualization and statistics about the weights of convolutional Layers

CNN takes the initial image in the input layer. The image information is processed by the following layers, which produce the final classification (deletion or non-deletion). DeepSV displays the two classifications and corresponding confidence values. Furthermore, DeepSV provides visualization and statistics about the weights and activations of each layer in the network. Figure S4 shows the process of classifying one testing image, from the initial image into the network, to the final recognition of the image results.

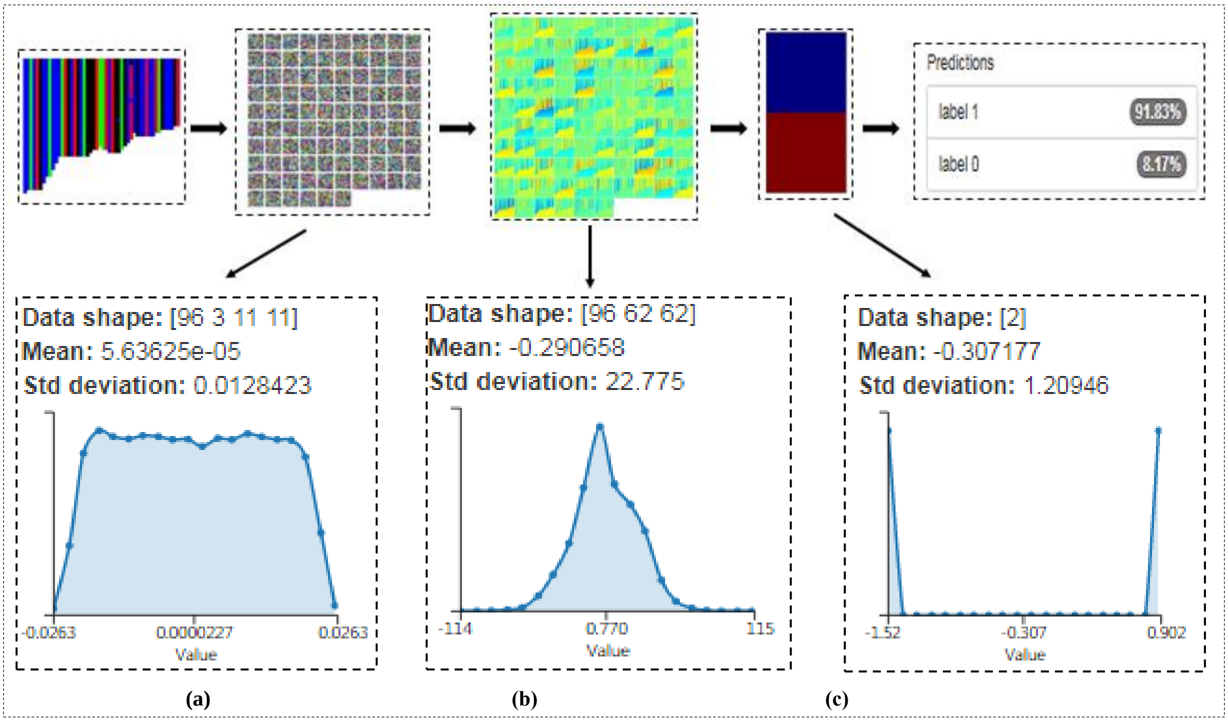


**Fig.S4.** How images are processed by layers in the CNN. Images are first processed by the input layer. Parts (a), (b) and (c) show the parameters of various convolutional layers at different stages. In the plots, horizontal axis: range of weight values; vertical axis: the number of weights. Data shape represents the size of the current feature map.

#### 3.5 PCA of Model's Fully Connected Layers

PCA (principal component analysis) is the method of projecting high-dimensional data into 3D space. We know that the last three layers of the DeepSV's CNN network are FC (fully connected layers). In order to verify the quality of model training, PCA was performed on the weights of the last three layers. The purpose of PCA is to maximize the mapped variance and maintain the relative distance between the original weight and the weight after mapping. The Figure S5 reflects the FC's PCA. It can be seen from the figure that the mapping is very dense, indicating that the weights are very dense and the model is trained well when most of these points after PCA are concentrated in 0. It is worth noting that the first and second layer are connected to the convolutional layer and the number of weights is huge. The last layer is connected to the softmax layer with relatively small number of weights.

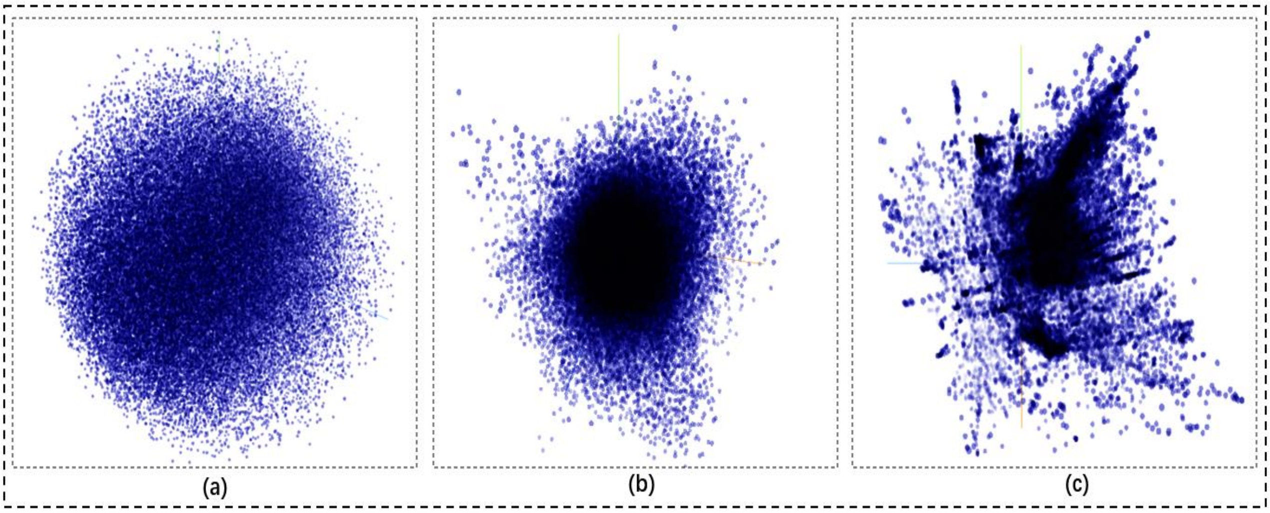


**Fig.S5.** The PCA result of fully connected layers. (a) the first full connected layer of PCA. (b) the second full connected layer of PCA. (c) the last full connected layer of PCA.

### 4 Picture labeling and normalization

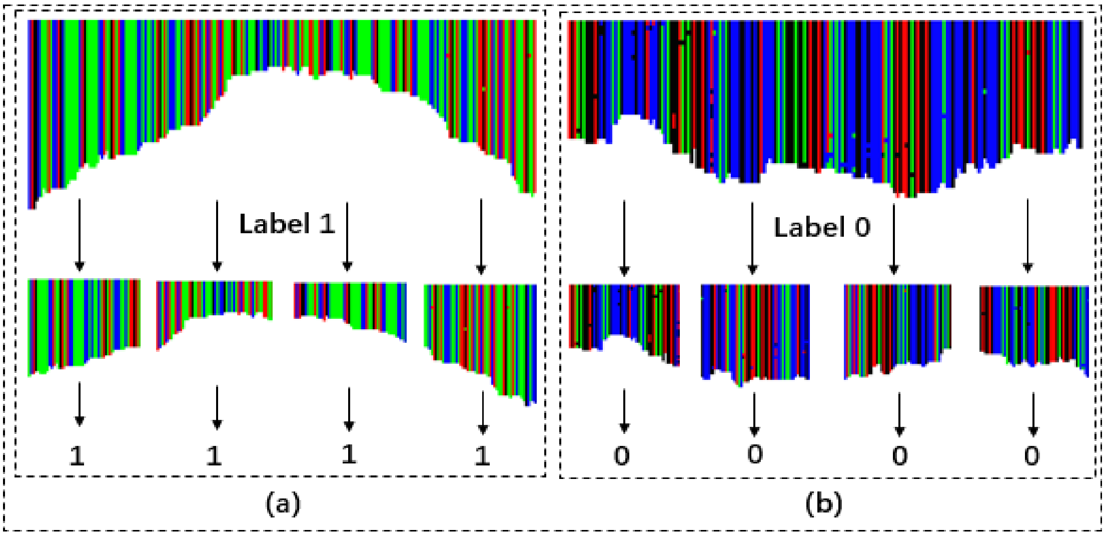


**Fig.S6.** (a) Deletions are divided into small fragments with label 1 which are sent into model training. (b) Non-deletions are divided into small fragments with label 0 which are also sent into model training. Each fragment is fixed length.

### 5 Deletion calling

Once the CNN model is trained, it is used for deletion calling on test data. To call deletions, DeepSV takes the aligned reads in the BAM format and reference genome. DeepSV generates pileup images in exactly the same way as in training. These images are given to the CNN model, which classifies the images to either deletions (1) or non-deletions (0). A contiguous sequence of 1s indicates the presence of a deletion. Here, some noises are allowed (e.g. one 0 in the middle of long sequence of 1s). The breakpoints of the deletion correspond to the first and the last 1s in this sequence. Note that breakpoints obtained this way may not be exact if the true breakpoints don't coincide with the image center. Breakpoints can be made more accurate by using the split reads as in Pindel. The called deletions are output in a VCF file that contains the specific breakpoints and the length of the deletion region. In addition, users can manually enter any two positions on the reference genome to detect whether there exists a deletion between the two positions. If there exist deletions within the two given positions, DeepSV finds the exact deletion regions. This process is illustrated in Figure S7.

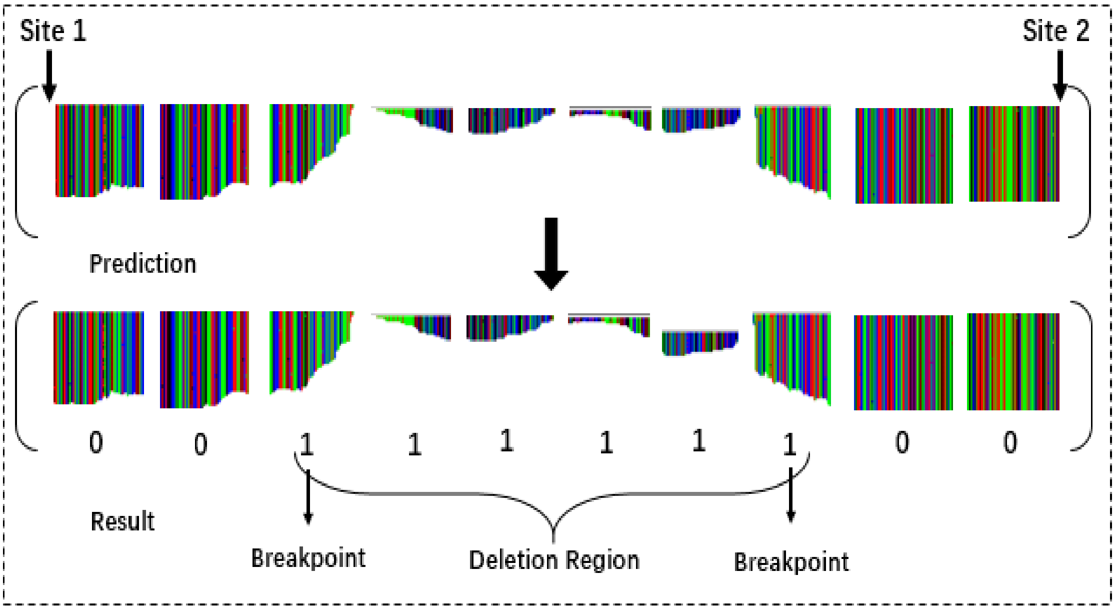


**Fig.S7.** The flow diagram of DeepSV's prediction. We send unknown pictures to DeepSV for detection and the DeepSV will tag each picture on testing results. From each image marked with a specific label, it can be based on the value of 0,1 to determine the deletion region and breakpoints.

### 6 Commands and Data used in the experiments

Commands used for running the tools, data used in the experiments.

#### 6.1 Command line options for running existing tools on deletion calling

(1). Pindel: pindel -f $ref -i ${sample}_${chr}${downsample}.pindel.cfg -o out.pindel.${sample}_${chr}${downsample} -c $chr -r false -t false -w 0.1 -x 5 -B 0 -T 4Sadf

(2). BreakDancer: bam2cfg.pl -q 20 ${sample}_${chr}${downsample}.bam > ${sample}_${chr}${downsample}.breakdancer.cfg && breakdancer-max ${sample}_${chr}${downsample}.breakdancer.cfg>out.breakdancer.${sample}_${chr}${downsample}

(3). DELLY: delly -t DEL -o out.delly.${sample}_${chr}${downsample} -g $ref ${sample}_${chr}${downsample}.bam

(4). CNVnator: cnvnator -genome ${ref} -root ${1}.root -chrom chr${2} -tree ${bamf}

(5). Breakseq2: run_breakseq2.py --bwa ${bwaDir} --samtools ${samtoolsDir} --bplib_gff ${gffFile} --bams ${bamFile} --chromosomes ${chr} --reference ${refFile} --work ${workDir} --nthreads ${threadNum}

(6). Lumpy: lumpy -mw 4 -tt 0 -pe id:${sampleName},bam_file:${lumpyPath}${sampleName}.discordants.bam,histo_file:${lumpyPath}${sampleName}.histo,mean:${mean_insertsize},stdev:${issd},read_length:${readLength},min_non_overlap:${readLength},discordant_z:5,back_distance:10,weight:1,min_mapping_threshold:20 -sr id:${sampleName},bam_file:${lumpyPath}${sampleName}.splitters.bam,back_distance:10,weight:1,min_mapping_threshold:20 > ${lumpyPath}${sampleName}_lumpyResult

(7). SVseq2: svseq -r $ref -b ${sample}_${chr}${downsample}.bam -c $chr --o out.svseq.${sample}_$chr

(8). GenomStrip2: SV_DIR=path_to_svtoolkit

classpath="${SV_DIR}/lib/SVToolkit.jar:${SV_DIR}/lib/gatk/GenomeAnalysisTK.jar:${SV_DIR}/lib/gatk/Queue.jar"

java -Xmx4g -cp ${classpath}

java -Xmx4g -cp ${classpath}

#### 6.2 List of individuals from the 1000 Genomes Project used for training and testing

**Table S3.** Individuals are used in the experiments

| Individuals Names |
| --- |
| NA12878 NA18511 NA18525 NA18631 NA18643 NA19017 NA19238 NA19239 NA19240 NA19625 NA19648 NA20502 NA20845 NA20868 NA20870 NA20885 NA20902 NA21090 NA21098 NA21125 |

### 7 The result of supplements

#### 7.1 The result of deletion calling of different deletion sizes on low coverage data

We compare the accuracy of deletion calling of different deletion sizes on DeepSV and other calling tools with low coverage data. To measure the performance of deletion calling, we use the following statistics: precision (P), sensitivity (S), and the F-score. The results on low coverage are shown in Table S4. The results on high coverage are shown in Table S5.

**Table S4.** Performance of deletion calling of different sizes on low coverage data from the 1000 Genomes Project (average coverage 10x). All the deletions are divided into different intervals according to the normal distribution. Various indicators are measured on detecting deletions with different tools. **P: precision S: sensitivity F: F-score = 2*P*S/(P+S).**

| Length Distribution | 50bp~200bp | | | 200bp~500bp | | | 500bp~1kbp | | | 1kbp~5kbp | | | 5kbp~10kbp | | |
| --- | --- | --- | --- | --- | --- | --- | --- | --- | --- | --- | --- | --- | --- | --- | --- |
| | P | S | F | P | S | F | P | S | F | P | S | F | P | S | F |
| Pindel | 0.29 | 0.61 | 0.39 | 0.57 | 0.64 | 0.60 | 0.68 | 0.34 | 0.45 | 0.52 | 0.37 | 0.43 | 0.11 | 0.12 | 0.11 |
| BreakDancer | 0.16 | 0.04 | 0.06 | 0.27 | 0.64 | 0.38 | 0.31 | 0.55 | 0.40 | 0.72 | 0.64 | 0.68 | 0.30 | 0.17 | 0.22 |
| Delly | 0.00 | 0.00 | 0.00 | 0.16 | 0.63 | 0.26 | 0.22 | 0.69 | 0.33 | 0.55 | 0.63 | 0.59 | 0.24 | 0.25 | 0.24 |
| CNVnator | 0.00 | 0.00 | 0.00 | 0.03 | 0.07 | 0.04 | 0.12 | 0.20 | 0.15 | 0.22 | 0.15 | 0.18 | 0.29 | 0.10 | 0.15 |
| Breakseq2 | 0.70 | 0.75 | 0.72 | 0.69 | 0.71 | 0.70 | 0.62 | 0.75 | 0.68 | 0.53 | 0.70 | 0.60 | 0.51 | 0.68 | 0.58 |
| Lumpy | 0.11 | 0.15 | 0.13 | 0.21 | 0.16 | 0.18 | 0.36 | 0.41 | 0.38 | 0.32 | 0.21 | 0.25 | 0.38 | 0.41 | 0.39 |
| GenomeStrip2 | 0.28 | 0.59 | 0.38 | 0.45 | 0.32 | 0.37 | 0.69 | 0.40 | 0.51 | 0.76 | 0.58 | 0.66 | 0.72 | 0.51 | 0.60 |
| SVseq2 | 0.25 | 0.52 | 0.34 | 0.57 | 0.76 | 0.65 | 0.80 | 0.41 | 0.54 | 0.65 | 0.42 | 0.51 | 0.26 | 0.11 | 0.15 |
| DeepSV | 0.81 | 0.73 | 0.77 | 0.77 | 0.63 | 0.69 | 0.72 | 0.76 | 0.74 | 0.57 | 0.65 | 0.61 | 0.52 | 0.51 | 0.51 |

From Table S4, we can see that Pindel and SVseq2 work better for medium size deletions with low coverage data. Delly and CNVnator don't perform well on deletions shorter than 200bp. The precision and sensitivity of BreakDancer, Lumpy and GenomeStrip2 increase as the deletion length increases. Breakseq2 has better sensitivity. Overall, DeepSV method performs well in all settings and outperforms the other tools in several settings (when deletion sizes are not too large) on low coverage data.

**Table S5.** Performance of Calling deletions of different sizes on high coverage data from the 1000 Genomes Project (average coverage 60x). Most tools have higher accuracy, sensitivity and F-score values than low coverage data. **P: precision S: sensitivity F: F-score = 2*P*S/(P+S).**

| Length Distribution | 50bp~200bp | | | 200bp~500bp | | | 500bp~1kbp | | | 1kbp~5kbp | | | 5kbp~10kbp | | |
| --- | --- | --- | --- | --- | --- | --- | --- | --- | --- | --- | --- | --- | --- | --- | --- |
| | P | S | F | P | S | F | P | S | F | P | S | F | P | S | F |
| Pindel | 0.32 | 0.33 | 0.32 | 0.51 | 0.43 | 0.47 | 0.42 | 0.48 | 0.45 | 0.62 | 0.44 | 0.51 | 0.21 | 0.25 | 0.23 |
| BreakDancer | 0.33 | 0.51 | 0.40 | 0.39 | 0.60 | 0.47 | 0.51 | 0.49 | 0.50 | 0.69 | 0.74 | 0.71 | 0.22 | 0.56 | 0.32 |
| Delly | 0.00 | 0.00 | 0.00 | 0.23 | 0.59 | 0.33 | 0.37 | 0.66 | 0..47 | 0.72 | 0.76 | 0.74 | 0.26 | 0.20 | 0.23 |
| CNVnator | 0.00 | 0.00 | 0.00 | 0.60 | 0.64 | 0.62 | 0.63 | 0.64 | 0.63 | 0.69 | 0.51 | 0.59 | 0.75 | 0.75 | 0.75 |
| Breakseq2 | 0.88 | 0.75 | 0.80 | 0.80 | 0.73 | 0.75 | 0.76 | 0.79 | 0.77 | 0.79 | 0.72 | 0.75 | 0.70 | 0.80 | 0.74 |
| Lumpy | 0.19 | 0.75 | 0.30 | 0.21 | 0.77 | 0.33 | 0.25 | 0.96 | 0.40 | 0.27 | 0.98 | 0.42 | 0.22 | 0.79 | 0.34 |
| GenomeStrip2 | 0.35 | 0.59 | 0.44 | 0.54 | 0.42 | 0.47 | 0.79 | 0.40 | 0.53 | 0.86 | 0.58 | 0.69 | 0.82 | 0.51 | 0.63 |
| SVseq2 | 0.22 | 0.59 | 0.32 | 0.47 | 0.62 | 0.53 | 0.71 | 0.30 | 0.42 | 0.55 | 0.40 | 0.46 | 0.16 | 0.13 | 0.14 |
| DeepSV | 0.97 | 0.95 | 0.96 | 0.94 | 0.95 | 0.94 | 0.90 | 0.92 | 0.91 | 0.85 | 0.83 | 0.84 | 0.65 | 0.75 | 0.70 |

As shown in Table S5, with high coverage of data, most tools perform better in precision and sensitivity than with low coverage data. For shorter deletions (less than 5 kbp), DeepSV tends to outperform the other tools on both precision and sensitivity. However, the precision and sensitivity of DeepSV on long deletions are slightly lower than those on shorter deletions. This is likely be due to the presence of complex variations for long deletions. For the shortest deletions (50bp~200bp), DeepSV achieves very high precision and sensitivity. In general, DeepSV performs well and outperforms other tools in almost all settings.

**7.2 Deletion calling for various frequencies**

**Table S6.** We divide the frequency into five intervals to measure the consistency of DeepSV. Sensitivity of calling deletions for deletions with different frequencies in population. Freq: number of individuals with a deletion among 20 individuals.

| Freq | Pindel | Break Dancer | Delly | CNVnator | Breakseq2 | Lumpy | Genome Strip2 | SVseq2 | DeepSV |
|---|---|---|---|---|---|---|---|---|---|
| **1-5** | 0.42 | 0.48 | 0.52 | 0.23 | 0.66 | 0.42 | 0.69 | 0.63 | 0.71 |
| **6-10** | 0.49 | 0.44 | 0.43 | 0.35 | 0.72 | 0.40 | 0.70 | 0.57 | 0.69 |
| **11-15** | 0.61 | 0.51 | 0.49 | 0.41 | 0.74 | 0.45 | 0.72 | 0.55 | 0.75 |
| **16-20** | 0.66 | 0.53 | 0.45 | 0.43 | 0.79 | 0.51 | 0.74 | 0.51 | 0.82 |

**7.3 Time and memory usage of different calling tools**

The running time and memory usage of DeepSV and other tools are shown in Figure S8. DeepSV is efficient in calling deletions. It is faster than other methods for deletion calling. Also the memory usage of DeepSV is also lower than most existing methods. Thus, the overhead for visualizing sequence reads appears to be low. We note that DeepSV takes relatively long time for model training (about 10 times more) than the time spent for deletion calling. With the training time included, DeepSV can be slower than some existing methods. Note, however, that model training is only performed once. The trained model can be used for calling deletions in many individuals.

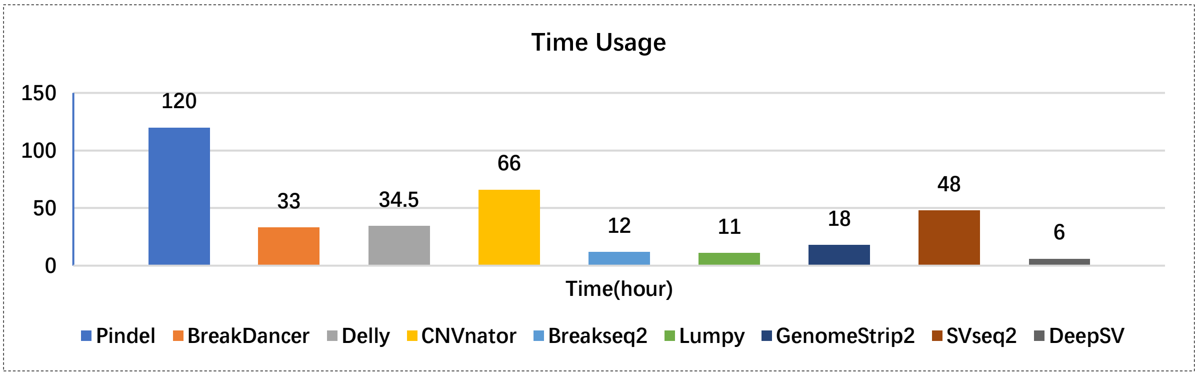


(a)

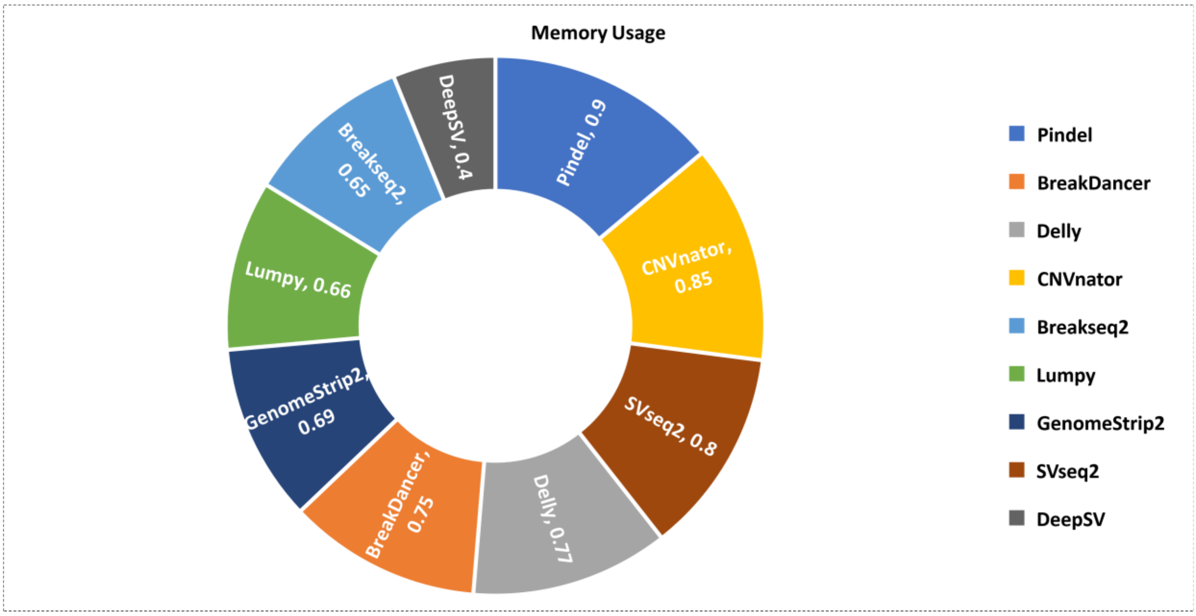

**(b)**

**Fig.S8.** Due to the different detection strategies used, the time spent by different tools to call deletions is different. Similarly, the amount of memory used is also different because of the amount of computation. (a) The prediction time of multiple tools. (b) The consumption of memory.
